# Molecular deconstruction of the pre-Bötzinger Complex/Nucleus Ambiguus (preBötC/NA) region: cellular constituencies and transcriptional responses to repeated seizures in the rat hindbrain

**DOI:** 10.1101/2025.09.22.677789

**Authors:** Wasif A. Osmani, Ibrahim Vazirabad, Neil Rhode, Gary Mouradian, Anna Manis, Matthew R. Hodges

## Abstract

Epilepsy affects millions worldwide, but a significant portion suffers from uncontrollable epilepsy. Repeated seizures have many consequences, including a high risk of post-ictal cardiorespiratory failure and Sudden Unexpected Death in Epilepsy (SUDEP). Major risk factors for SUDEP include biological sex in addition to the occurrence of generalized tonic-clonic seizures (GTCSs). How repeated seizures lead to cardiorespiratory dysfunction remains unknown. A key factor in many neurological diseases is neuroinflammation, predominantly mediated by microglia and astrocytes that become dysfunctional. Mechanistically, questions remain how they affect neuronal function in epilepsy and contribute to cardiorespiratory dysfunction and increased SUDEP risk. Previously, we have shown that repeated seizures in our novel rat model with genetic mutations in *kcnj16*, an inwardly rectifying K^+^ channel, in the Dahl salt sensitive rat (SS*^kcnj16-/-^*) led to increased neuroinflammation in key ventilatory regions at 3 and 5 days of seizures. Specifically, there was increased recruitment of various inflammatory mediators, increased recruitment of activated microglia, with improvement in post-ictal respiratory dysfunction and mortality with usage of anti-inflammatory agents. Here we tested the hypothesis that repeated seizures lead to differential neuroinflammatory activation after repeated seizures in CNS regions of ventilatory control. Male SS*^kcnj16-/-^* rats were subjected to 0 (Naïve), 3, 7 or 10 days of seizure, and subsequently, the pre-Bötzinger Complex/Nucleus Ambiguus (preBötC/NA) was isolated and sent for nuclei isolation and sequencing. Seurat was utilized to filter and process the data, integrate across conditions and allow for differential gene expression (DEG) analysis. Afterwards, pathways enrichment analysis was performed allowing for determination of unique pathways recruited across cell types for each seizure condition. Overall, we were able to identify 18 unique cell types based on transcriptomic signatures, with 8 different neuronal populations, grouped based on Type 1, Type 2 or a mixed Type 1 & Type 2 genetic expression, indicating rhythm generation or pattern generation, respectively. We found that majority of the neuronal clusters were Type 1 or mixed type, indicating predominantly rhythmogenic neuronal populations. Importantly, these critical neuronal populations showed significant upregulation in various metabolic and neurological disease pathways at the 3 and 7 Day timepoints. Furthermore, we identified various glial cells, including microglia and astrocytes and saw increased recruitment in various Inflammatory pathways, Metabolic pathways and Chemokine related pathways after 3 and 7Days of seizures, confirming our previous results. Consequently, our results show for the first time, transcriptomic characterization of crucial rhythmogenic neuronal populations after repeated seizures and the changes that may underlie their dysfunction in SUDEP, mediated in part through the network change in upregulated inflammatory pathways in surrounding glial cells.

## Introduction

Epilepsy affects millions of people worldwide and presents a large health-related issue for many caregivers around the world. A portion of these individuals suffer from uncontrollable epilepsy, putting them at risk for cognitive issues, but chiefly, Sudden Unexpected Death in Epilepsy Patients (SUDEP) (1). SUDEP is a clinical diagnosis of exclusion, and unfortunately disproportionately affects those suffering from uncontrollable epilepsy, male gender, and those who experience a higher frequency of generalized tonic-clonic seizures (GTCSs) (1,2). MORTEMUS, a landmark study, was performed by researchers to understand clinically the pathophysiology of this phenomenon (3). Using clinical epilepsy monitoring units (EMUs), they observed a stereotypical pattern in those that succumbed to SUDEP: a GTCS led to decreased breathing, followed decreased heart rate and then terminal apnea and asystole (3). This led to an important step in determining how cardiorespiratory centers in the brainstem may be negatively affected by seizures.

Neuronal networks that govern cardiorespiratory control reside in the brainstem. Some key brainstem nuclei include the nucleus of the solitary tract (NTS), which serves as an integration site of afferent information from the periphery sensing information from the carotid bodies and relaying that information to surrounding nuclei to modulate output of breathing and heart rate (reviewed in 4). Also, the nucleus ambiguous (NA), which serves to provide motor output to muscles of airway tone and regulate parasympathetic tone of the heart. Lastly, the pre-Bötzinger Complex (preBötC) serves as the central rhythm and pattern generator for breathing, providing signals to premotor neurons of the hypoglossal nuclei and phrenic to coordinate airway tone and downstream muscles to inflate the lungs for inspiration. It also receives inputs from surrounding regions that respond to changes in metabolic demand in the body, allowing fine control of breathing to meet changes in pH, O_2_ and CO_2_ (reviewed in 4). Importantly, this nucleus performs these crucial functions due to housing key neuronal subpopulations that are responsible for its rhythmogenic activity (5). Therefore, to understand the effect of repeated seizures on cardiorespiratory function, we chose to assess the preBötC/NA region from a cellular and transcriptional level.

The pre-Bötzinger complex (preBötC) houses critical populations of neurons that serve as the central rhythm and pattern generating centers in the brainstem (Reviewed in 4). Various groups of neurons in this medullary region have projections to other critical cardiorespiratory control centers. Studies have been able to determine markers for these groups of neurons, dividing them into inhibitory or excitatory interneurons with roles in different aspects of ventilatory control (6, 7). Due to the ventilatory changes seen in our audiogenic seizure model and patient data for those who succumbed to epilepsy, strong rationale exists to determine what may be happening in the preBötC after repeated seizures in a cell type specific manner to understand cell specific effects in these key neuronal control populations. Therefore, we performed snRNA seq to perturb this question, hypothesizing that key neuronal populations such as *tac1* (tachykinin, precursor 1), GABAergic and glutamatergic neurons, undergo detrimental transcriptomic changes after repeated seizures.

## Methods

### Animals and Tissue isolation

Age matched males (8-16 weeks) were selected for all experiments. Animals were maintained on a low salt diet and were only used for experiments once they were at least 8 weeks of age. Animals were placed in plethysmographs used for physiology measurements and underwent exposure to auditory tone for 2 min to elicit a level 3 or 4 seizure (8, 9). Between 60-90 minutes afterwards, on the designated time point for extraction, animals were euthanized using isoflurane and brains extracted and flash-frozen using ice cold isopentane. Brains were then mounted and sectioned using a cryostat to advance to Bregma locations for identified regions of interest (Identify obex (occurs at Bregma -13.56mm) Figure 146 in Rat Brain atlas (**Figure 1**). Once at obex, advance 6 x 120 um thick sections to get to Bregma -12.84mm Figure 140 in Rat Brain atlas for preBötC/NA, then 3x200 um punches were made using a steel rod with 0.5 mm diameter. Samples were stored in RNAase free tubes at -80C until they were sent to Singulomics (New York City, New York) to perform the nuclei isolation and 10X genomics sequencing.

**Figure 1.**
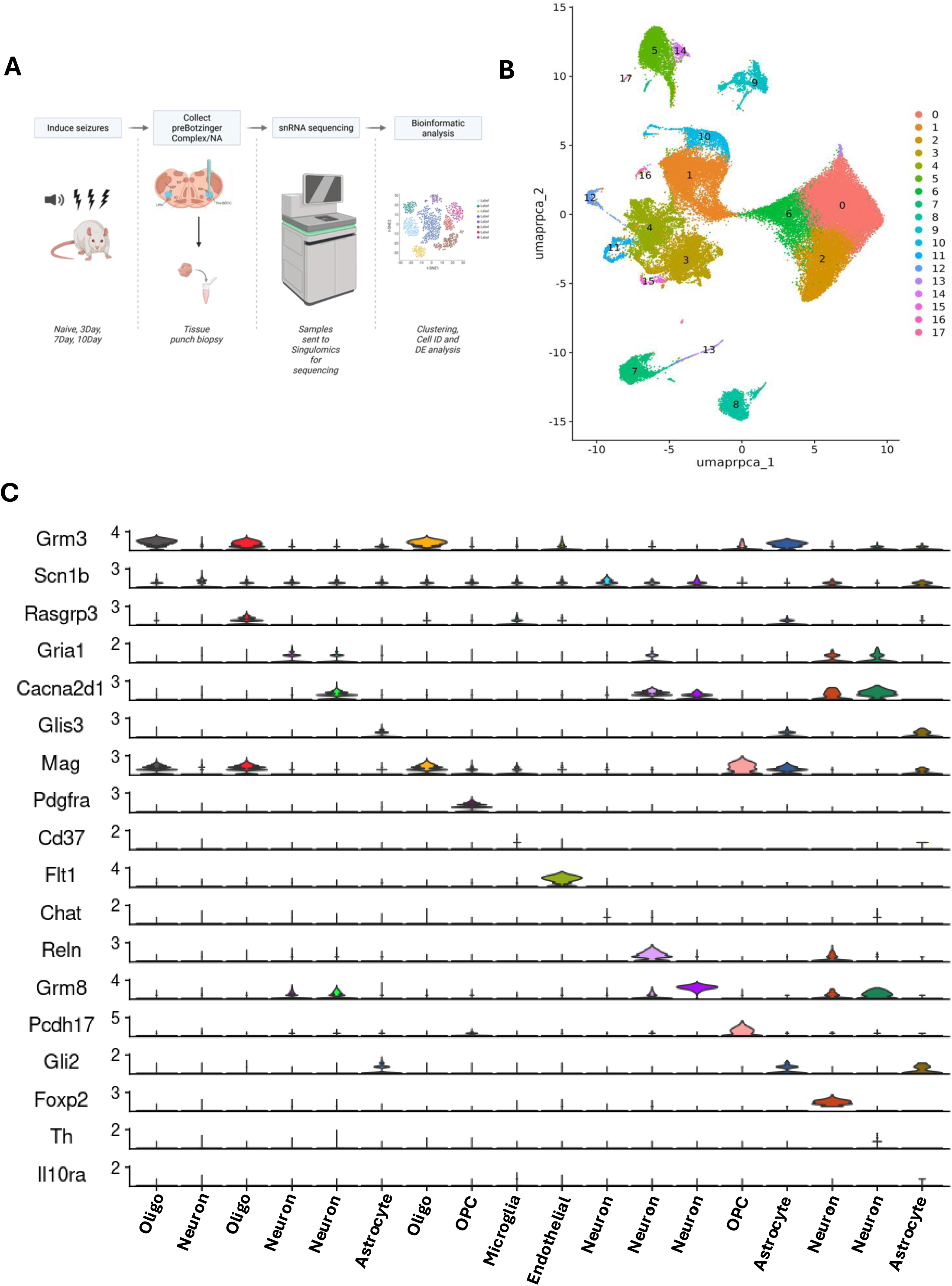
Seizure induction and single nuclear (sn)-RNA sequencing of preBotzinger Complex/Nucleus Ambigous (preBotC/NA). Methodology of seizure induction of male SS*^kcnj16-/-^* rats, isolating preBotC/NA, and samples sent for sn-RNA sequencing (**A**). **B** shows the integrated UMAP highlighting the 18 unique cell types following processing, integration and clustering of cells. **C** highlights through a violin plot, the most highly differentially expressed genes, using Seurat’s FindMarkers function, that allow characterization of cell type. PangloDB was subsequently used to identify cell type based on the top 10 highly expressed genes for each cluster, of which the most unique gene is plotted here for the subsequent cluster, denoting a unique cell type.

### Integrating sequence data from multiple timepoints

An initial group of males (n =1/group) was sequenced, targeting 5,000 nuclei per sample, with 2 runs of HiSeq lane of 400 million PE150 reads. Clean reads were then analyzed with the rat reference genome Rattus_norvegicus 6.0.97 using Cell Ranger v3.1.0 (performed by Singulomics). Across the four samples, there was an average of 10,850 ± 1,024 cells, 20,250 ± 2,092 reads/cell, 1,002 ± 90 genes/cell, and 213,324,217 ± 237,070 reads per sample. Additionally, quality control scores (Q30) for different measures were as reported across the 4 samples (average ± SEM): Q30 Bases in Barcode = 97.73% ± 0.03%, Q30 Bases in RNA Read = 88.78% ± 0.14%, Q30 Bases in UMI = 97.75% ± 0.03%. With all quality scores for sequencing reporting in Q30 values, with at least near 90% accuracy, this shows that the reads are of high quality with very little probability of inaccuracy. Also, across the 4 samples, there was a 91.75% ± 0.43% of reads that mapped to the genome, 80.13% ± 0.65% of reads that mapped confidently to the genome, 59.38% ± 1.23% of reads mapped confidently to exons, and 32.13% ± 2.50% of reads mapping confidently to the transcriptome. After this initial sequencing run, a second round of animals (n = 2) were added to increase power to give a total of 3 animals for each condition. Following the same procedures and methods as the initial run, samples were collected and sent to Singulomics for processing and sequencing. Again, 5,000 nuclei were targeted utilizing a NovaSeq 6000 (PE150) for sequencing, clean reads were analyzed with rat reference genome Rattus norvegicus 6.0.97 using Cell Ranger v6.0.1 (performed by Singulomics). On average, across the samples, there was an average of 7,563 ± 1,042 cells captured 32,723 ± 4,770 reads/cell, 861 ± 50 genes/cell, and 234,556,828 ± 16,996,287 reads per sample. Quality control scores (Q30) scores across the samples for this run were as follows (average ± SEM): Q30 Bases in Barcode = 95.90% ± 0.27%, Q30 Bases in RNA Read = 88.53% ± 1.35%, Q30 Bases in UMI = 94.55% ± 0.59%. Also, across the samples, there was a 82.93% ± 3.58% of reads that mapped to the genome, 71.00% ± 3.42% of reads mapped confidently to the genome, 29.58% ± 2.62% reads that mapped confidently to exons, and 32.23% ± 3.72% reads that mapped confidently to the transcriptome. Overall, the reads generated from these samples were of high quality and mapped to the proper regions in a similar manner to the previous samples. Therefore, this enabled us to confidently integrate samples to generate a larger dataset to work with that included 3 animals per condition.

### Filtering sequence data, clustering and UMAP generation

To process the sequence datasets, we utilized the Seurat v3 and updated v5 guided tutorial as the basis of our standardized pipeline to process the raw data we received from Singulomics to filter out mitochondrial DNA (5% cutoff), to scale and normalize the data accordingly for the individual timepoints e.g. naïve, 3Day, 7Day, and 10Day, and determine cutoffs for variable features for each individual timepoint before integrating via SCT in Seurat (6, 10). Integration was performed using a modified version of the Seurat Integration vignette. Once the individual timepoints were created into SeuratObjects, scaled, and filtered, they were integrated using CCA (Canonical correlation analysis) for dimensional reduction (1:50). Once integration was completed, the integrated SeuratObject was scaled and principal component analysis (PCA) performed using 50 PCs (*p <* 0.05 for PCs). PCA was utilized for dimensionality reduction to generate a UMAP with the Euclidean metric with 1:50 dimensions. FindNeighbors, utilizing a PCA reduction and 1:50 dimensions, was used to determine the nearest neighbors within the integrated dataset. Lastly, FindClusters was used with a resolution of 0.34 and the standard Louvain algorithm to generate clusters of defined, nearest neighbors. scCustomize v3 was utilized for visualization of plots and to display data across timepoints utilizing its built-in features for multiple displays.

## Results

### Clustering, UMAP and Cell-type identification

Using the methods noted earlier, we were able to identify 18 unique cell populations in the NA/preBötC region (**Figure 1B**). Due to the large variety of neuronal subtypes within this region, we desired an unbiased and objective approach to determine cell type prior to determining the individual subtypes of neuronal populations. Therefore, we generated a function that utilizes the top 10 expressing genes based on log fold values, for each cluster to have a list of genes acting as markers to help identify that unique cluster. FindAllMarkers was utilized to generate gene markers and expression values. The function utilizes PangloDB, a single cell sequence database that houses user sequence datasets that can be used to determine where and what types of cell express a given gene or gene list (11). After inputting the Top 10 expressed genes for each cluster into the database, we had the cell type identities for the integrated SeuratObject: Endothelial cell – Cluster 9; Oligodendrocyte – Clusters 0, 2, 6; Astrocyte – Cluster 5, 14, 17; Oligodendrocyte Precursor Cell (OPC) – Clusters 7, 13; Microglia – Cluster 8; Neurons – Clusters 1, 3, 4, 10-12, 15, 16. **Figure 1C** shows the top expressed marker gene for each cell type and its expression for the respective cell type, as visualized by a violin plot. Therefore, confirming the cell type that was determined using PangloDB. Given that we were able to identify various cell types present in the NA/preBötC region, we next sought to determine the identifies of the critical neuronal populations present. To confirm physiological and functional relevance of the neuronal types identified, we sought to determine if we could find unique, gene markers that may represent important neuronal populations that have been identified in the literature and in various studies, amongst the neuronal clusters (6, 7, 12). Based on work done by various groups, there have been key neuronal subpopulations of crucial relevance that can roughly be categorized into interneurons, excitatory and inhibitory neurons (6, 7, 12). Recent work by Del Negro et al characterized the inspiratory neurons through single cell sequencing of the preBötC from neonatal mice, establishing various electrophysiological markers of key ion channels, proteins and markers that contribute to either rhythm or pattern generation (7). Moreover, their classification of Type 1 and Type 2 neurons, Type 1 being rhythmogenic and Type 2 being pattern-generating was applied here to provide a framework in delineating the unique neuronal subtypes observed in this dataset. Utilizing the markers Del Negro et al highlighted, we selected markers for both Type 1 and Type 2 neurons, separated by the broad category of electrophysiological nature of the gene (**Figure 2**). Type 1 electrophysiological markers *kcnj9, kcnk1, kcna4,* and *scn1b* encode potassium ion channel subunits and Na channel subunits that contribute to the A-type transient K+ current (I_A_), whereas the non-electrophysiological markers *gria2, grin1, gabra3, glrb*, *penk* and *oprl1* encode ionotropic, metabotropic, and neuropeptide subunits for GABAergic and glycinergic inhibitory neurons (**Figure 2B, D**). Type 2 neurons express hyperpolarization-activated mixed cationic current (I_h_), dependent on electrophysiological markers such as *kcnn3, kcnt2, kcna6, scn8a,* and *cacna2d3*, representing various potassium ion channel and calcium channel subunits (**Figure 2C**). Non-electrophysiological markers such as *gabra1, grik5, agt, sst, tacr3, mcoln1,* and *trpm7* represent various GABA, kainate, peptide and TRP-channel subunits that contribute to I_h_ (**Figure 2D**). Our dataset contained predominantly a mix of Type 1 non-electrophysiological marker expressing neurons and Type 2 electrophysiological neurons, seen in all clusters except for cluster 10, which appeared to express predominantly Type 1 markers specifically (**Figure 2B-E**). Given the diverse array of expression of the markers in many clusters, each cluster may contain neurons of the different types shown, contributing to the expression on the violin plots. Consequently, this approach to characterizing the various neuronal clusters seen in our dataset confirmed the presence of various crucial rhythmogenic and pattern-generating neuronal populations. Once the various neuronal populations were identified based on key marker expression into Type 1, Type 2 or mixed type, a final UMAP (**Figure 3**) was generated to portray the various cell types and their markers used to identify them. Importantly, key Choline acetyl transferase (*Chat*^+^) neurons were also identified confirming that initial tissue punches of preBötC/NA were accurate, as *Chat* serves as an important landmark gene marker for the nucleus ambiguous (NA).

**Figure 2.**
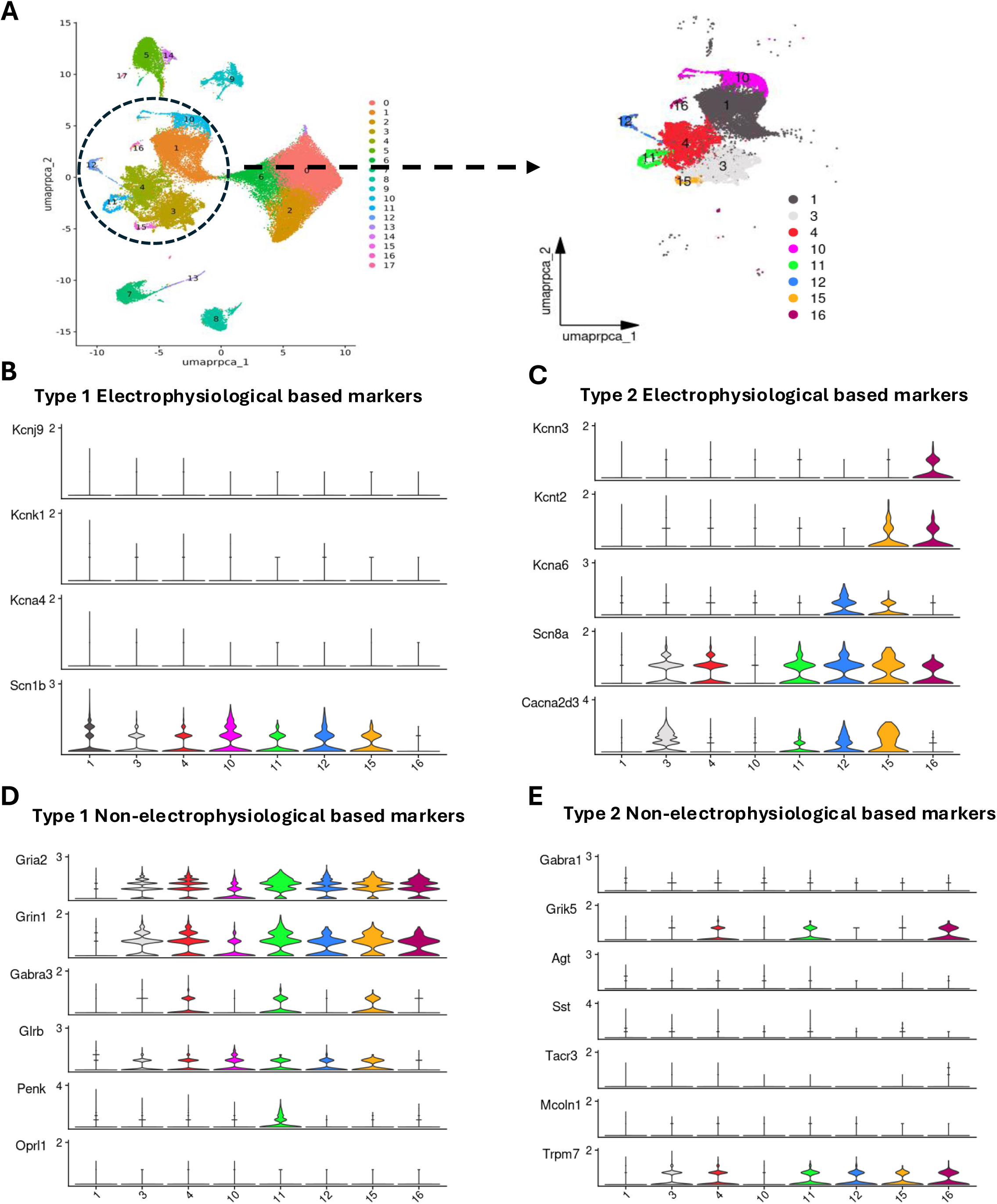
Neuronal classification based on gene expression parameters. **A** represents the focus on the clusters 1, 3-4, 10-12, 15-16 as the neuronal clusters based on the PangloDB classification based on the top 10 highly expressed genes. **B-E** highlights the methodology used to further characterize and classify the neuronal clusters into unique groups, based on gene expression profiles for Type 1 and Type 2 neurons. **B** and **D** highlights violin plots showcasing the expression of key genes based on electrophysiological markers for Type 1 (**B**) or non-electrophysiological markers in Type 1 neurons (**D**). **C** and **E** highlights violin plots showcasing the expression of key genes based on electrophysiological markers for Type 2 (**C**) or non-electrophysiological markers in Type 2 neurons (**E**).

**Figure 3.**
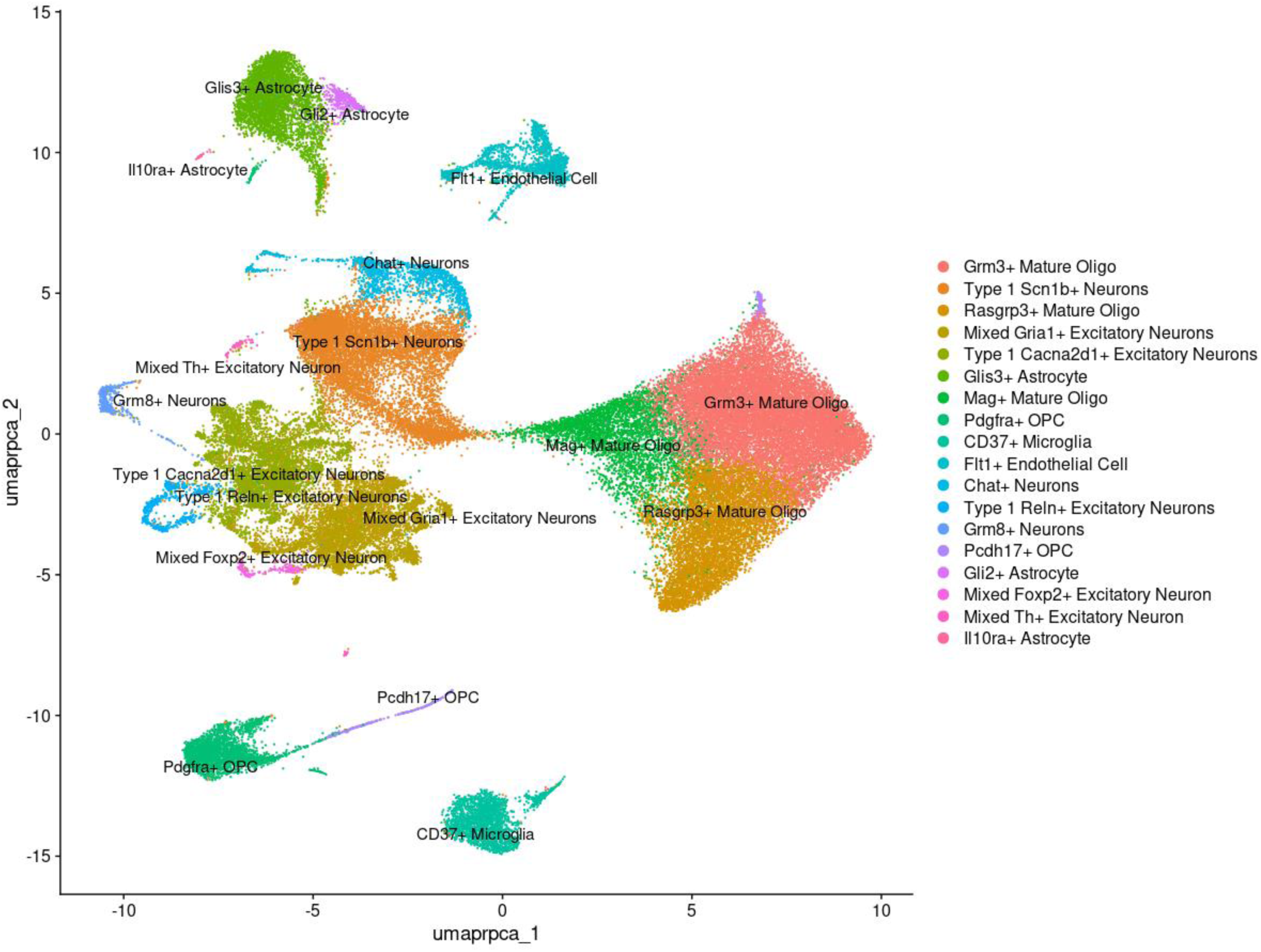
Identified cell types after sn-RNA sequencing of preBotC/NA. This UMAP represents the finalized Seurat object after identifying cluster cell types utilizing FindMarkers and PangloDB, and identifying specifically Type 1 and Type 2 neurons, based on key gene expression markers shown previously.

### Differences in Gene expression after repeated seizures

After defining key rhythmogenic populations of neurons and defining unique cell types, we sought to determine what the effect of repeated seizures may be on gene expression in these important neuronal populations and the region. Therefore, we utilized Seurat’s FindMarkers package to compare the naïve condition with the seizure timepoints and determine which genes were differentially expressed in the seizure condition relative to naïve. Differentially expressed genes (DEGs) were generated for those that had an adjusted *p*-value < 0.05 and for each individual cluster. Initial observations (**Figure 4A**) showed that across seizure timepoints, there were varying changes in the density of rhythmogenic preBötC clusters. To determine the effect on differential gene expression, we sought to see how many unique DEGs were expressed at each seizure timepoint (**Figure 4B**). Using a triple Venn diagram, we see that the 3Day timepoint had the greatest number of unique DEGs (1029), followed by 7Day (798) and then 10Day (367). This suggests that the greatest change to the system occurred within the earlier timepoint, after which, the system was able to adapt with more seizures, leading to less DEGs with more days of seizures. Importantly, we saw a core group of 832 DEGs that were consistent across the repeated seizures, suggesting that these were key genes that were consistently perturbed and may play an important role in regulating the system’s response to repeated insult. To more closely assess these core genes, a heatmap was generated to determine any directional change of the top 26 most significant core genes (**Figure 4C**). Similarly, the greatest change in expression occurred at the 3Day timepoint, with most genes showing increased expression relative to naïve, followed by some genes showing a switch from downregulation at 3Day to upregulation at 10Day (*rpl9, ppia, cox6a1*) (**Figure 4C**). Additionally, volcano plots were generated to determine which genes for each time point had the greatest change, relative to naïve, either upregulated or downregulated (**Figure 4D-F**). Again, the greatest change in up- and downregulated genes was present in the 3Day timepoint (**Figure 4D**). Of note, the genes identified as the core top 26 genes (**Figure 4C**) represent genes prevalent in various energy metabolism and ribosomal processing functions, suggesting repeated seizures targeting key functions of crucial cell types of the region.

**Figure 4.**
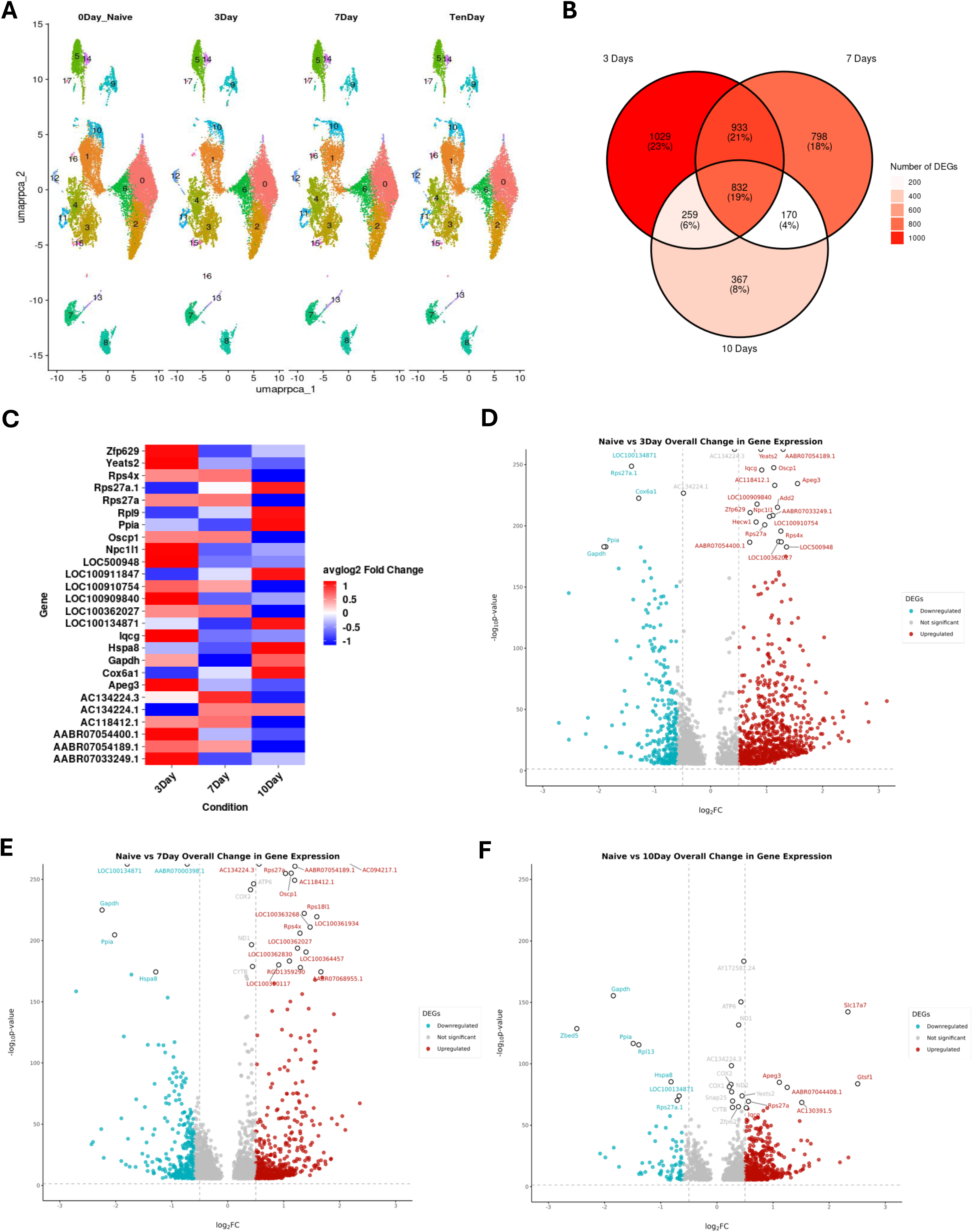
Overall changes in gene expression after repeated seizures. **A** shows the changes in UMAP cluster density over time with repeated seizures. **B** shows a triple Venn diagram of the DEGs of the 3Day, 7Day and 10Day condition, relative to naïve, and the genes that are unique and common across the conditions, with the darker a color (darker red gradient) indicating increasing number of DEGs for that condition. **C** represents the top 26 most significant (*p* < 0.05) common DEGs across all the 3 conditions, plotted in a heat map with their average logarithmic fold change expressed on a color gradient with blue indicating decreased expression relative to naïve and red indicating increased expression relative to naïve. **D-F** shows volcano plots of each condition and the DEGs, with blue indicating down regulated and red indicating up regulated relative to naïve, with threshold log2FoldChange = 0.5, and with genes plotted higher on the y-axis, indicating increased significance (increasingly small *p*-value).

### Enrichment and Pathway Analysis

After seeing broad changes in DEGs in the preBötC/NA after repeated seizures, we sought to understand what pathways may be changing because of the DEGs in specific cell types. Therefore, we used enrichment analysis (ShinyGO) to perform a pathway analysis from the DEG lists for neurons, microglia, astrocytes and endothelial cells across the seizure timepoints (13). Due to the large nature of the dataset and results that were generated for each cell type, results presented focus on the top 10 most significant pathways. Therefore, a select few cell types of physiological and functional relevance will be highlighted in this section.

### Neurons

As noted earlier, we were able to identify various Type 1 and Type 2 neuronal populations within our dataset, along with mixed type neurons that express markers for both rhythmogenic and pattern generating function (**Figure 3**). To assess how repeated seizures were affecting these crucial neuronal populations, DEGs were generated as before (**Figure 5A, B**) and interestingly, the greatest number of DEGs were present at the 7Day timepoint (2707), then the 10Day timepoint (614), and lastly by the 3Day timepoint (263). This suggests that the initial insult of 3 Days of seizures may not perturb the system as significantly to cause any major detrimental changes, however, continued repeated insults may have a cumulative effect that reaches its zenith at the 7Day timepoint. Additionally, when looking at the core DEGs similar across all three timepoints, there were 1782 genes that were differentially expressed, suggesting that these genes may play a crucial role in adapting to insults and maintaining homeostasis from stressors to ensure that key functions remain intact. We plotted the top 25 most significant core genes and saw that within these genes; the most significant change was upregulation at the 3Day timepoint relative to naïve (**Figure 5B**). Interestingly, 21 of the 25 most significant core genes were upregulated, however, 4 genes were downregulated, specifically *calm1, loc100134871, cox6a1, rps27a.1* (**Figure 5B**). To determine the function of the core 25 genes, we utilized a chord diagram and plotted the KEGG pathways associated with the genes to show which genes were clustering towards unique pathways (**Figure 5C**). Importantly, majority of the genes were associated with the Ribosomal pathway, followed by the Metabolic pathways, then Parkinson Disease pathway, suggesting that the core neuronal DEGs function predominantly in Ribosomal, Metabolic and neurologic disease pathways. Similarly, we sought to determine which pathways were enriched after the different seizure timepoints (**Figure 5D-F**). Beginning with the 3Day timepoint, we saw Oxidative Phosphorylation as the most highly enriched and one of the most significant pathways, followed by Ribosome pathway (**Figure 5D**). The remaining top 10 enriched pathways were represented by pathways related to various neurological disease or disease processes, such as Pathways of neurodegeneration-multiple diseases, all of which were positively enriched, predictive of upregulation. Consequently, this suggests that at the 3Day timepoint, the neuronal populations undergo vast insult leading to upregulation of various disease associated pathways putting strain on the rhythmogenic neuronal populations while trying to meet energy and metabolic needs. At the 7Day timepoint, Lysosome and Protein processing in the ER were among the most enriched KEGG pathways identified, however, of the top 10 pathways, half of them were associated with various neurological disease pathways e.g. Amyotrophic lateral sclerosis, Parkinson Disease and Pathways of neurodegeneration-multiple diseases. All pathways presented in the plot at the 7Day timepoint were upregulated, indicating activation because of 7Days of seizures. Similar to the 3Day timepoint, pathways related to metabolic function, Metabolic pathways, was the most significant pathway identified at 7Day, suggesting that along with the continued stress to the system and damage response through various neurological disease related pathways, high metabolic demand remains to allow for the system to respond. At the 10Day timepoint, there appeared to be a clear and distinct change in the pathways enriched, as majority of the pathways were related to axonal guidance, synapse maintenance and protein processing, with the most enriched pathway being Synaptic vesicle cycle. Of note, Pathways of neurodegeneration-multiple diseases was enriched at the 10Day timepoint still, suggesting continued positive activation of damage response to the system. However, with the majority of enriched pathways resembling synaptic maintenance and inhibitory synapse activity, the 10Day timepoint appears to show a system that has adapted to the repeated seizures, with a focus on developing and building new and remaining synapses (**Figure 5F**).

**Figure 5.**
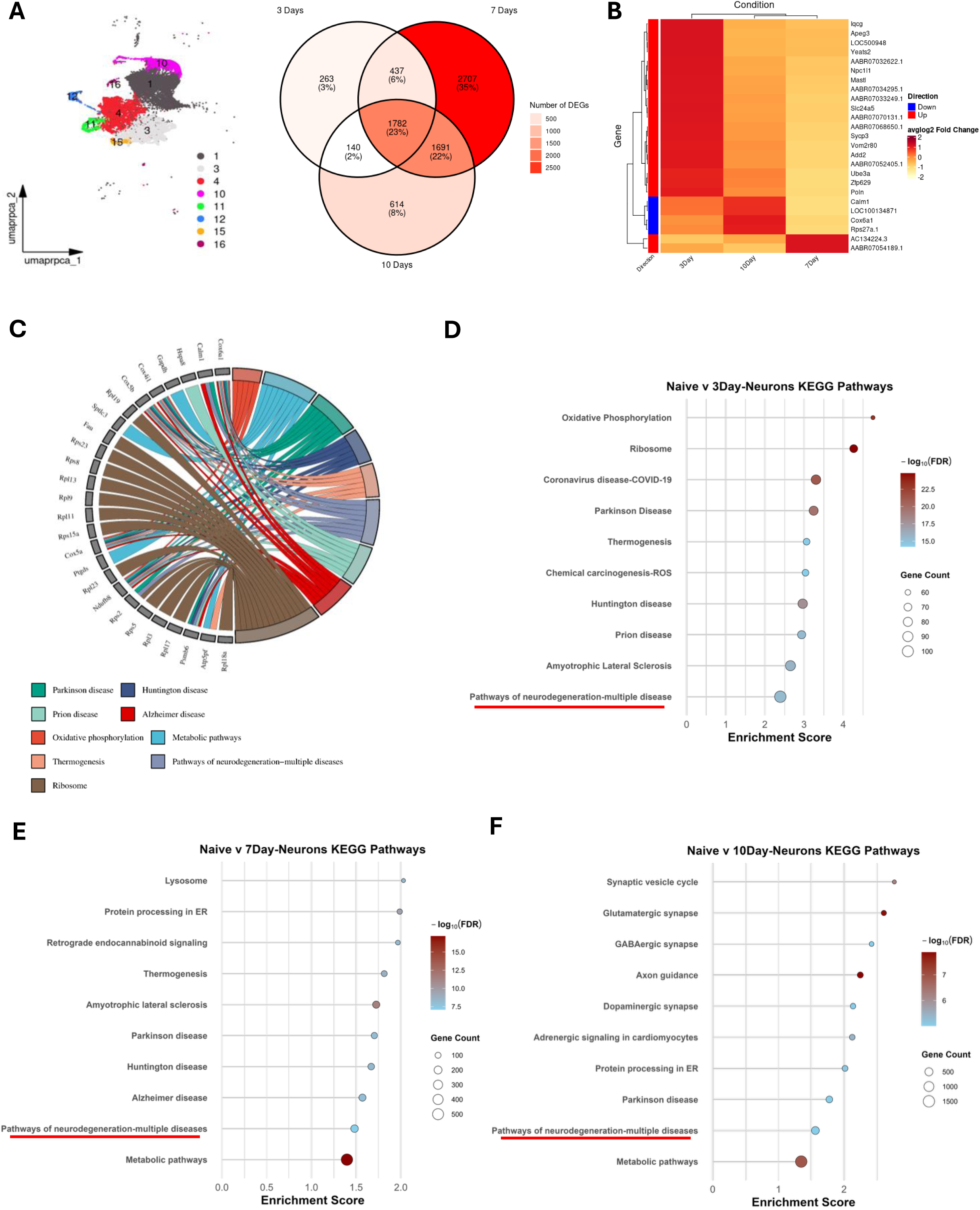
Neuronal differential gene expression and pathway analysis after repeated seizures. **A** shows the neuronal clusters that were grouped together to allow for a more robust analysis for DEGs and GO enrichment pathway analysis, showing in a triple Venn Diagram the common DEGs across each condition, relative to naïve. The darker red color indicates an increasing number of DEGs. **B** shows the top 25 most significant common DEGs across all 3 conditions, plotted in a heat map with color indicating increased expression (the darker red indicates higher avglog2Fold Change) with inclusion of directional changes relative to naïve plotted with blue color denoting down regulation. Conditions were hierarchically clustered based on similarity of DEG changes for that condition. **C** represents a GO Chord diagram plotting the top 25 most common DEGs and the GO terms that they were enriched in, showcasing the core pathways the genes were involved in. **D-F** plots the top 10 KEGG enriched pathways for each condition, with enrichment score plotted on the x axis indicating greater positive enrichment or upregulation, color change of blue to red indicating greater significance (smaller FDR < 0.05) and size of circle denoting the number of DEGs included. Red lines under pathways highlight common neurodegenerative pathways included in all three conditions.

### Glial cells and Endothelial Cell effects

Utilizing a similar approach to characterize and assess pathway changes in neuronal groups, we investigated effect of repeated seizures on microglia, astrocytes and endothelia cells. Glial cells, both microglia and astrocytes, have various roles in CNS immune regulation and maintaining homeostasis in regulating neuronal function, especially after insult (14–17). Previously, we have shown increased numbers of microglia after repeated seizures in our model and shown that there is a concomitant increase in the expression of various chemokines and cytokines, in a site and time dependent manner, correlating with morphological changes in microglia within this key rhythmogenic region (9). Therefore, assessing microglia, astrocytes and endothelial cells, which have been shown to have various roles in regulating CNS immune function, may help elucidate the impact repeated seizures have on the region. **Figures 6** and **7** show overall, the top 10 most significant (*p* < 0.05) pathways that were affected across the seizure timepoints, focusing on KEGG pathways and Gene Ontological (GO) Biological and Cellular pathways that revealed significant changes of physiological and functional relevance. Within the CD37^+^ microglial population, relative to the naïve condition, the most significant pathway that was upregulated following 3 days of seizure, was *B cell Receptor signaling* (-log(FDR) > 5.5, Enrichment score = 14), followed by *Fc Gamma receptor-mediated phagocytosis* (-log(FDR) >5.5, Enrichment score = 12). Among the top 5 most significant and upregulated pathways was *Chemokine signaling pathway* (-log(FDR) > 5.5, Enrichment score = 8). Importantly, this supports our previously published findings showing increased chemokines and cytokines after 3 and 5 days of seizure, along with the increase in number and morphological changes that reflected an activated glial state, characterized by increased release of inflammatory mediators (9). Similarly, at the 7 Day timepoint (**Figure 6B**), the top 10 pathways collectively reflect pathways involved in various neurological disease states and infections. Specifically, the second most significant and positively enriched pathway was *Coronavirus disease-COVID19* (-log(FDR) > 6, Enrichment score = 8). Among the top 10 most significantly enriched pathways was *Pathways of neurodegeneration-multiple diseases* (-log(FDR) > 3, Enrichment score = 3.25), along with *Parkinson’s Disease, Prion Disease, Huntington’s Disease and Alzheimer’s Disease,* all positively enriched, indicating these pathways were significantly upregulated following 7 Days of seizure. At the 7Day timepoint, the upregulated pathways all reflected disease pathways that were similarly seen in the neuronal populations, suggesting damage mediated possibly by the microglial cells as the sentinel immune cells of the CNS. At the 10Day timepoint (**Figure 6C**), the most significantly upregulated pathway was *Glycolysis/Gluconeogenesis* (- log(FDR) > 3.5, Enrichment score = 16), followed by different metabolic related pathways. However, *Pathways of neurodegeneration-multiple diseases*, *Parkinson Disease, Prion Disease,* were among the top 10 pathways as well, suggesting that various disease processes were still prevalent at 10Days of repeated seizures, however, metabolic shifts reflecting different energy requirements may imply that microglial activity shifted to adapt to a chronically inflamed state.

**Figure 6.**
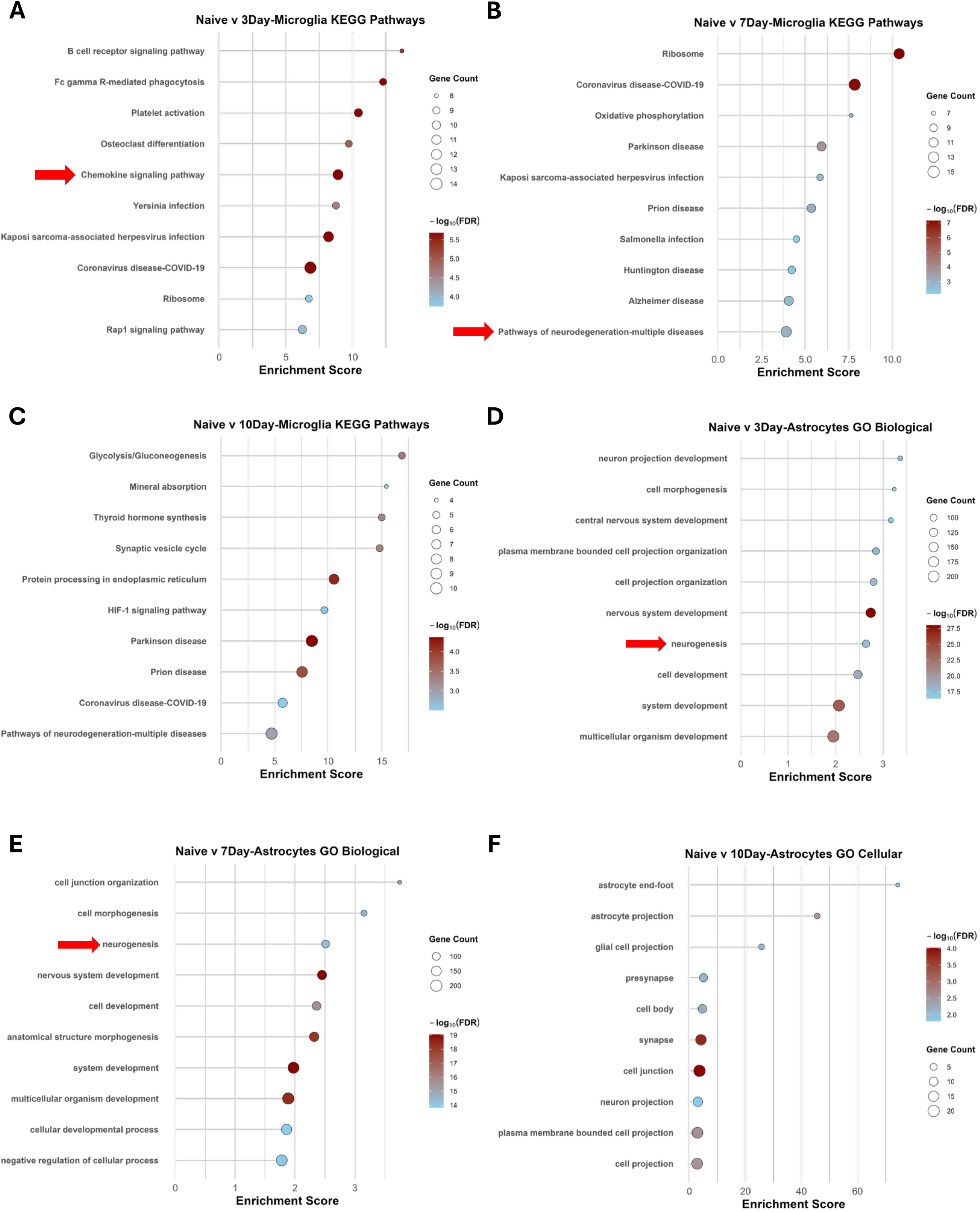
Microglial and Astrocyte Enrichment Pathway analysis after repeated seizures. **A-C** plots the top 10 KEGG enriched pathways for each condition for microglia, with enrichment score plotted on the x axis indicating greater positive enrichment or upregulation, color change of blue to red indicating greater significance (smaller FDR < 0.05) and size of circle denoting the number of DEGs included. **D-F** plots the top 10 GO enriched pathways for each condition for astrocytes, with enrichment score plotted on the x axis indicating greater positive enrichment or upregulation, color change of blue to red indicating greater significance (smaller FDR < 0.05) and size of circle denoting the number of DEGs included. Red arrows highlight key pathways that were upregulated and involved immune/inflammatory relevance.

**Figure 7.**
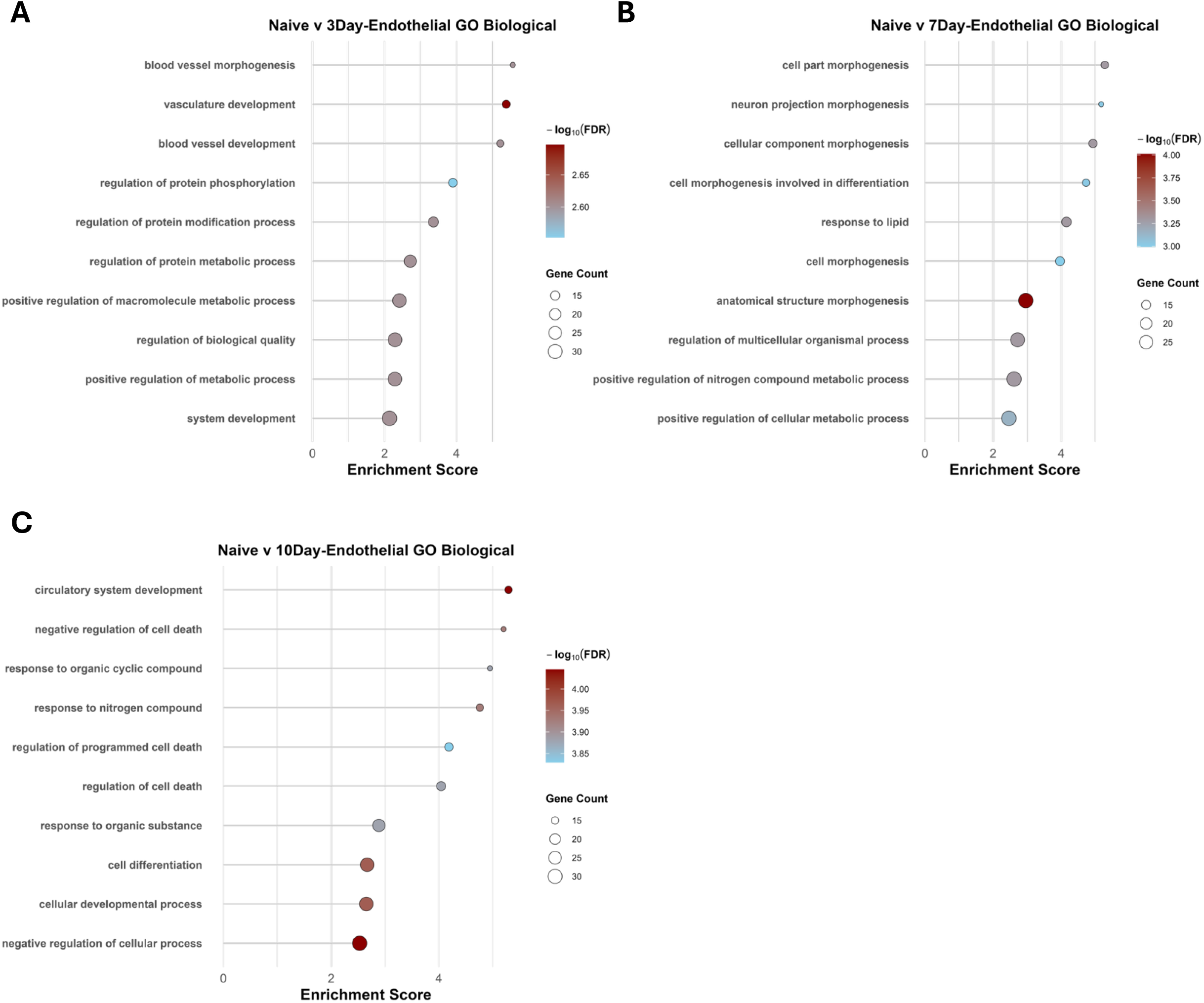
Endothelial cell enrichment pathway analysis after repeated seizures. **A-C** plots the top 10 GO enriched pathways for each condition for endothelial cells, with enrichment score plotted on the x axis indicating greater positive enrichment or upregulation, color change of blue to red indicating greater significance (smaller FDR < 0.05) and size of circle denoting the number of DEGs included.

Our sequencing analysis resulted in identifying 3 unique astrocytic clusters, identified by markers, Glis3^+^, Gli2^+^ and IL10ra^+^ astrocytes (**Figure 3**). For the enrichment analysis, similar to the neuronal enrichment analysis, all three clusters were grouped to allow for increased *n* in assessing pathway changes given IL10ra^+^ diminished density relative to the other 2 astrocytic populations. As done with the neuronal and microglial populations, enrichment analysis was performed on the grouped astrocytic population to assess for which pathways astrocytes in our dataset were changed after repeated seizures. **Figure 6 D-F** shows the representative changes for the top 10 most significant pathways. **Figure 6D** shows the top 10 most significant GO Biological pathways that were significantly upregulated relative to the naïve condition after 3 days of repeated seizures. Within the astrocytes, at 3Days of repeated seizures, the most significantly upregulated pathways reflect overall changes in neuronal development and cellular morphogenesis, suggesting astrocytic involvement in restoration of surrounding neuronal architecture. Moreover, the most enriched pathway was *Neuron projection development* (-log(FDR) >17.5, Enrichment score = 3.4), with the most significantly upregulated pathway being *Nervous system development* (-log(FDR) > 27.5, Enrichment score = 2.7). Among the top 10 most significantly enriched pathways, *Neurogenesis* (-log(FDR) >17.5, Enrichment score = 2.6) was the 7^th^ most significantly upregulated enriched pathway, suggesting one of most important function of the astrocyte after 3Days of seizure was to upregulate genes that support neurogenesis in the region. At 7Days of seizure (**Figure 6E**), the top 10 most significantly enriched upregulated pathways again focused on neuronal development, morphogenesis, cellular development and neurogenesis. The most highly enriched pathway was *Cell junction organization* (-log(FDR) > 14, Enrichment score = 3.8), while the most significantly upregulated pathway was *Nervous system development* (-log(FDR) > 19, Enrichment score = 2.45). *Neurogenesis* pathway was noted as the 3^rd^ highest enriched pathway at 7Days of seizure (-log (FDR) > 14, Enrichment score = 2.5), suggesting after 7Days of repeated seizures, the system as whole displays a greater need for regrowth and neurogenesis to continue to function. At the 10Day timepoint, utilizing the differentially expressed genes, enrichment analysis investigating pathway changes using GO Biological changes did not reveal significant pathways, therefore, GO Cellular changes were assessed, which revealed various cellular processes within the astrocyte population, such as synapse function, astrocyte end foot function and cell projection (**Figure 6F**). The most significantly upregulated and enriched pathway noted was the *Astrocyte end foot* pathway (-log(FDR) > 2, Enrichment score = 75), and the most significant pathway was noted to be *cell junction* pathway (-log(FDR) > 4.0, Enrichment score = 4), followed closely by the *synapse* pathway (-log(FDR) >4.0, Enrichment score = 5). Overall, at the 10Day timepoint, astrocytic cellular pathway involvement suggested further recruitment of morphological changes aimed to support further growth of the surrounding architecture via increased synapse and end-foot development. Lastly, our analysis investigated endothelial cells, a cell type that has been shown to play an increasingly important role in the preservation of the blood brain barrier, allowing for the infiltration of various peripheral immune cells and the diffusion of various cytokines and chemokines in neuroinflammatory processes (reviewed in 18). Therefore, we sought to understand, what role if any, do endothelial cells play after repeated seizures through characterizing the transcriptomic changes in this key region. **Figure 3** revealed the identification of a single endothelial cell cluster in our dataset, characterized by *Flt1+* expression. **Figure 7** revealed the pathway enrichment analysis, similar to the neuronal and glial cells, of the top 10 most significant pathways following repeated seizures in the endothelial cell population. **Figure 7A** shows that the most significant pathway after 3Days of seizures was the upregulation of *blood vessel morphogenesis* followed by *vasculature development* (-log(FDR) > 2.6, Enrichment score = 5.5; -log(FDR) > 2.65, Enrichment score = 5.3). Overall, the pathways that were most significantly upregulated at the 3Day seizure timepoint were pathways involved in various metabolic and regulatory processes (**Figure 7A**). **Figure 7B** reveals that after 7Days of seizures, the top 10 most significantly upregulated pathways focused on various morphological changes to the system, with the most significantly enriched pathway being *cell part morphogenesis* (-log(FDR) > 3.25, Enrichment score = 5.2). Given the change in the enriched pathways at the 7Day seizure timepoint compared to the 3Day timepoint, this suggests a switch in focus in the endothelial cells responding to greater levels of insult to the system *i.e.* seizures and adapting through structural and morphological changes. At the 10Day seizure timepoint (**Figure 7C**), the most significantly enriched pathway was the *circulatory system development* (-log(FDR) > 4, Enrichment score = 5.2), followed by *negative regulation of cell death* (-log(FDR) >3.90, Enrichment score = 5.15). Of note, included in the top 10 pathways was *regulation of programmed cell death* (-log(FDR) > 3.8, Enrichment score = 4.1) and *regulation of cell death* (-log(FDR) > 3.85, Enrichment score = 4), numbered 5 and 6, respectively. Importantly, with the inclusion of cell death pathways as enriched at the 10Day timepoint, this suggests that endothelial cells play a role in regulating the turnover of nearby cells, potentially cells associated in the blood-brain barrier.

## Discussion

The novelty of this experimental approach resides in being able to isolate a key cardiorespiratory control region and identify various cell types and subpopulations of neurons that have contributed to preBötC function (7). We show here in the identification of various neuronal groups that express markers that have been shown to be expressed in unique rhythm-generating neurons. Within the males, there were both excitatory (*slc17a6*^+^ - *vglut2*) and inhibitory (*slc32a1*^+^ - GABAergic) neurons, along with those that can be putatively described as Type 1 preBötC neurons (related to a I_A_ current – A type transient K^+^ current and are responsible for the pre-inspiratory summation of synaptic potentials that lead to the inspiratory burst and are rhythmogenic) and Type 2 preBötC neurons (express a I_H_ current – a hyperpolarization-activated cation current and activate ∼300 ms after Type 1 neurons, can be understood as pattern-generating) based on their expression of channels that contribute to the respective currents (7). *Sst^+^* expression was more closely associated with Type 2 neurons, whereas *cacna2d1^+^* (calcium channel) was associated with Type 1 neurons. A larger group that comprises a heterogenous group of preBötC neurons are those expressing *scn1b* (voltage gated sodium channel subunit) and these neurons that expressed along with a *gria2* (excitatory glutamate channel subunit), which likely represent a unique population that includes rhythmogenic and pattern-generating neurons, defined as a mixed-type. Interestingly, our results show that the large predominant types of neuronal populations that were identified, majority were Type 1 or a mixed-type, with no sole Type 2 neuronal population isolated based on our analysis.

Within this context, transcriptomic changes resulting from seizures in these neurons suggest major time dependent changes in function. Mounting evidence shows the recruitment of inflammation in a variety of neurological disorders, including epilepsy, and so using DEGs we sought to determine what changes may be occurring in different cell types and important neuronal populations (14–15, 18–19). There were clear changes in neuronal clusters in recruitment of various pathways that were immune related or showed oxidative phosphorylation to be significantly affected, as evidenced by the GO Chord diagram using the core genes that were consistently differentially expressed across seizures (**Figure 5C**). Due to the high metabolic demands of neurons in general, decreased energy production and dysfunctional mitochondria may underlie poor neuronal health (20–21). By not meeting the necessary demands of the cell, it can affect the postsynaptic output and potential from that neuron, creating the potential for dysfunction in inspiratory rhythmogenesis (20–21). Moreover, Friese et al showed that in a rodent model of multiple sclerosis, mitochondrial dysfunction was prevalent, and by overexpressing a mitochondrial transcriptional co-regulator in neurons, there was improved neuronal function, while deleting it led to worsened neurodegeneration (22). This suggests another line of evidence supporting the idea for mitochondrial function playing an important role in affecting neuronal function, especially when assessed in a neurological diseased state.

Over the 10 days, the neuronal populations continued to show changes in metabolic pathways and a recruitment of immune and inflammatory related pathways, upregulated to confirm immune involvement post-seizure, even after 10 days of seizures (**Figure 5F**). Overall, there appears a link in inflammation occurring after repeated seizures, driven by activated microglia and astrocytes that may be causing neuronal dysfunction and changes in energy metabolism, negatively affecting their function, consistent with our published data in Osmani et al 2024 (9). Specifically, we showed time and site dependent recruitment of microglia, with a significant increase in inflammatory mediators, with the greatest increase after 5 Days of seizures. Given that the 5 Day timepoint was not a part of this experimental approach due to animal constraints, utilizing the DEG and pathway analysis at the 3Day and 7Day timepoint, we saw recruitment of various inflammatory pathways, corroborating our previously published study. Therefore, this indicates that neuronal dysfunction, along with inflammatory recruitment, rather than neuronal death could compromise vital functions, such as breathing and cardiovascular function. Future studies will confirm the presence of these markers and changes in gene expression through protein analysis and explore how mitochondrial dysfunction may be intertwined with inflammation.

